# Purifying circular RNA by ultrafiltration

**DOI:** 10.1101/2024.12.04.626383

**Authors:** Karen Guillen-Cuevas, Xiaoming Lu, Marc R. Birtwistle, Scott M. Husson

## Abstract

Developing messenger RNA (mRNA) vaccines for COVID-19 renewed and intensified the interest in using mRNA for disease prevention and treatment. Despite their efficacy, linear mRNA molecules are short-lived in the human body, primarily due to enzymatic degradation at the free ends. In contrast, circular RNA (circRNA) exhibits enhanced stability and resistance to exonuclease degradation. However, this stability depends highly on purity. Unfortunately, the *in vitro* transcription/self-splicing reaction products contain a mixture of circular and linear RNAs. Yet, practical methods for purifying circRNA from solutions containing linear RNA contaminants are lacking. In this study, we explored the feasibility of using ultrafiltration to purify protein-encoding circRNA produced by the self-splicing of a precursor RNA (preRNA) during *in vitro* transcription (IVT). We measured the sieving coefficients, a separation metric, of circRNA, linear precursor RNA, and nicked RNA conformers using polyethersulfone membranes with molecular weight cutoffs from 30 to 300 kDa, analyzing performance as a function of permeate flux. We also estimated the RNA critical fluxes and determined suitable operating conditions for purification. We achieved a purity of 86% with a yield above 50%. By comparison, the purity achieved by size-exclusion high-performance liquid chromatography (SE-HPLC), the leading alterative separation technology, was 41% with a yield of 45%. These findings highlight ultrafiltration as a superior method for purifying circRNA at the research scale. Its scalability suggests that it could play a critical role in enabling the large-scale manufacturing of circRNA-based therapeutics.

## Introduction

Messenger RNA (mRNA) therapies reached an important milestone in 2020 when the U.S. FDA issued the first medical authorization for an mRNA vaccine to prevent COVID-19. The news coverage on the effectiveness of the Pfizer/BioNTech and Moderna vaccines significantly increased societal awareness of mRNA and its ability to deliver the coding sequence for therapeutic protein production to the body to prevent disease. Since then, mRNA therapeutics have taken a leading role in developing new drugs and vaccines[1]. Despite the high efficacy of the COVID-19 vaccines, the linear structure of the encapsulated mRNA molecules makes them innately susceptible to enzymatic degradation at the free ends, which shortens their half-life *in vivo* and limits the duration of therapeutic protein expression. Therefore, additional steps must be taken to reduce the innate cellular immune response and protect mRNA from degradation. These include nucleoside modifications, codon optimization, and the addition of expensive synthetic 5’ cap analogs, which have yielded only modest improvements in stability[2, 3].

Circularizing RNA eliminates the free ends associated with exonuclease-mediated degradation. Circular RNA (circRNA) is well-established for translation and can be protein-coding[2, 4, 5]. Thus, circularizing protein-coding RNA is a promising strategy for increasing the stability, duration, and quantity of therapeutic protein production[3, 6–8]. For example, circRNA cancer vaccines show greater stability than linear mRNA vaccines[9].

The primary strategy to produce engineered circRNA is to incorporate self-splicing introns. In this method, a linear mRNA precursor (preRNA) is designed with 3’ and 5’ introns that self-splice through transesterification reactions, resulting in circularization. [4]. Wesselhoeft et al. [3] used such a system to engineer circRNA up to 5 kb in length with high circularization efficiency and demonstrated stable protein production when delivered to eukaryotic and mammalian cell lines. Subsequently, they showed that purified circRNA decreased immune responses upon cellular entry relative to unpurified circRNA, which contains linear RNA and byproducts, making them an attractive vector for therapeutic gene expression[3]. Thus, overall, the shape of circRNA may lead to better, longer-lasting therapies than linear mRNA.

In addition to the potential therapeutic benefits associated with their improved stability circRNA plays roles all across biology and disease[9]. They arise from back-splicing pathways thought to be ubiquitously present in most human cells. They are expressed in all human tissues[8, 9]. To better understand the roles circRNA plays across such diverse biology, tools to obtain high yields of pure circRNA will be necessary for biomedical research.

For engineering applications, circRNA is synthesized using an *in vitro* transcription (IVT) back-splicing reaction. The circRNA synthesis produces solutions containing the plasmid DNA template, residual preRNA, nicked (i.e., damaged) RNA, and circRNA. The protein expression stability of circRNA depends strongly on purity, with small quantities of the contaminating preRNA and nicked RNA leading to robust cellular immune responses[3]. Therefore, circRNA must be purified to remove these and other contaminants.

Methods to purify circRNA include gel electrophoresis, exonuclease degradation and size exclusion chromatography[10]. Gel electrophoresis is used to separate the RNA conformers (preRNA, nicked RNA, and circRNA), which exhibit different migration rates. Although the three conformers are similar in molecular weight, their different secondary structures influence their migration rates during gel electrophoresis. Hence, the bands representing preRNA, nicked RNA, and circRNA are distinguishable. It is established that circRNA migrates slower than equal-weight nicked RNA in EX agarose E-gels[11]. Following electrophoresis, purified fractions of the RNA conformers can be obtained by gel extraction. Unfortunately, the heating during separation may compromise RNA stability and reduce the yield. Furthermore, electrophoresis methods are not scalable.

Exonuclease degradation can be used to enrich circRNA. This approach uses RNase R exonuclease to break down linear RNA. This method takes advantage of circRNA stability and resistance to exonucleases. However, circRNA may become nicked and degrade at purification conditions, especially at high RNase R concentrations. This enzyme’s high cost compared to the yield makes it unsuitable for mass production[10].

Size exclusion (SE) high-performance chromatography (HPLC) separates RNA molecules based on molecular weight and polarity differences. SE-HPLC is a good option for removing introns. However, the peaks overlap when separating RNA conformers whose sizes are highly similar. The selectivity is low. Thus, stringent selection of a narrow peak fraction is required to obtain high purity, causing low recovery yield. By combining SE-HPLC and RNase R enrichment, Wesselhoeft et al.[12] obtained a purity of 90% but with a yield below 50%. In addition to being ineffective for bench-scale purifications, these methods are considered nonviable options for commercial scale therapeutic circRNA purification[10].

Thus, developing new circRNA therapeutics requires novel scalable separation approaches to reach high purity while producing a yield high enough to make it suitable for mass production. This study explored whether ultrafiltration (UF) could purify circRNA from IVT reaction solutions with high purity and yield. UF has been used for plasmid DNA purification, where it has been shown that linear DNA has lower critical flux for transmission through a UF membrane than supercoiled and open circular isoforms[13]. We theorized that circRNA and linear (precursor and nicked) RNA similarly would have different critical fluxes. We hypothesized that if circRNA, preRNA, and nicked RNA have different critical fluxes, UF can separate them by carefully controlling the flux during diafiltration runs using an appropriate UF membrane.

## Materials and Methods

### circRNA synthesis and gel separation

A bacterial plasmid to transcribe circRNA encoding NanoLuc^®^ (circRNA-synIRES-R25-NanoLuc) was a gift from Dr. Howard Chang (Addgene plasmid #188116, Watertown, MA, USA)[2]. It was streak-plated in a kanamycin agar plate and cultivated for 18 h at 37 °C. The kanamycin agar plates were made using LB agar (Miller) (Fisher Bioreagents, Suwanee, GA, USA) and 50 µg/mL kanamycin sulfate (Apexbio Technology LLC, Houston, TX, USA). Individual colonies were selected and cultivated for 18 h at 37 °C in kanamycin LB media. The kanamycin LB media was made using LB Broth (Miller) (Fisher Bioreagents) with 50 µg/mL kanamycin sulfate (Apexbio Technology LLC). A ZymoPURE Plasmid Miniprep Kit (Zymo Research, Fisher Scientific, Suwanee, GA, USA) was used to extract the plasmid from the bacterial broth. The plasmid was directly amplified using Q5 polymerase using Polymerase chain reaction (PCR) (Q5® High-Fidelity PCR Kit, New England Biolabs, Ipswich, MA, USA). PCR was carried out following manufacturer protocol (**Table S3**) at a primer annealing temperature of 68°C for 30 cycles using a Bio-Rad C1000 Touch Thermal Cycler (Bio-Rad, Hercules, CA, USA). The reverse and forward primer sequences were provided by Chen et al.[14] and procured from Integrated DNA Technologies (Coralville, IA, USA).

circRNA was synthesized by IVT using the HiScribe T7 High Yield RNA Synthesis Kit (New England Biolabs, Ipswich, MA, USA) and the circRNA-synIRES-R25-NanoLuc DNA templates. One microgram of DNA template was used per 20 µL of IVT reaction volume. Reactions were run for 16 h at 37 °C. IVT templates were degraded with 2 µL of DNase I (New England Biolabs) per IVT reaction for 20 min at 37 °C. The remaining RNA was purified using a Monarch RNA Cleanup Kit (New England Biolabs). To probe circularization, purified RNA was treated with RNase R (Applied Biological Materials, Richmond, BC, Canada). RNase R is a 3′ to 5′ exoribonuclease that digests linear structures, but not circular ones.

IVT reaction products were diluted to approximately 20 ng/µL, mixed with 2X RNA loading dye (New England Biolabs) in a 1:1 v/v ratio, and heated at 70 °C for 10 min to denature the RNA. A molecular weight ladder was prepared using 2 µL of ssRNA ladder (New England Biolabs), 8 µL of RNase-free water (Thermo Fisher Scientific, Waltham, MA, USA) and 10 µL of 1X RNA loading dye. The ladder was denatured with the IVT samples. After denaturing, IVT samples and the RNA ladder were loaded into an Invitrogen E-Gel EX precast agarose gel (2%) (Thermo Fisher Scientific) at 20 µL per well and run on the E-Gel Power Snap Electrophoresis System (Catalog number G8300, Thermo Fisher Scientific) using the setting ‘‘EX-2%’’ for 15 min at room temperature. Images were taken using Bio-Rad ChemiDoc XRS+ and Image Lab 6.1 software using the “SYBR-GOLD’’ setting.

Purified fractions were obtained by gel RNA extraction. Gel electrophoresis separated circRNA, precursor RNA (preRNA), and nicked RNA. A spatula was used to open the E-Gel case. While using a blue light lamp and wearing orange safety glasses, a sterile razor blade was used to cut the gel fractions containing the circular, preRNA and nicked RNA. A Zymoclean Gel RNA Recovery Kit (Zymo Research, Thermo Fisher Scientific) was used to melt the agarose gel and extract the purified RNA fractions. The purified fractions were eluted in 50 µL of RNase-free water and stored at -80 °C.

### Characterizing RNA size by dynamic light scattering

The hydrodynamic diameters of the RNA conformers were measured using dynamic light scattering (DLS) with a Zetasizer (Malvern Panalytical, Malvern, UK) and an Ultra-Micro Cell for Nano Series quartz cuvette (ZEN2112, Malvern Panalytical). The sample volume was 15 µL, and the concentration was around 50 ng/µL.

### Ultrafiltration system

The system was based on Latulippe and Zydney[13, 15]. It consisted of a 50 mL Amicon stirred cell (UFSC05001, MilliporeSigma, Burlington, MA, USA) connected to an Amicon stirred cell reservoir (6028, MilliporeSigma) pressurized using compressed air. The pressure was regulated using a Wilkerson compressed air regulator (Grainger, Greenville, SC, USA) and measured using an Omega pressure transducer (Omega Engineering, Norwalk, CT, USA). The cell was stirred at 200 rpm. The cell volume was reduced to 10 mL by adding an insert fabricated in Clemson University Machining and Technical Services because the purified RNA fractions are laborious and expensive to synthesize (**Figure S1**). A 300, 100, 50 or 30 kDa nominal molecular weight cut-off (MWCO) Biomax^®^ polyethersulfone ultrafiltration membrane disc (Millipore-Sigma) was washed by soaking it for 16 h in diethylpyrocarbonate (DEPC)-treated water (Thermo Fisher Scientific) on a Fisherbrand Multiplatform Shaker (Thermo Fisher Scientific) and then loaded at the base of the cell. Permeate flow was adjusted by changing the exposed surface area of the membrane using waterproof aluminum tape.

### Permeability measurements

The membrane hydraulic permeability is the slope of the best-fit line on a plot of the permeate flux and the transmembrane pressure. The permeate flux was calculated as the volumetric flow rate through the membrane divided by the exposed surface area. The volumetric flow rate was measured gravimetrically using an Adventurer analytical balance (OHAUS Corporation, Parsippany, NJ, USA). The membranes were preconditioned by flowing 100 mL of DEPC-treated water at the highest pressure (345 kPa). The pressure varied from 69 kPa to 345 kPa with 69 kPa increments. Three measurements were taken at each pressure in the order of decreasing pressure and then repeated in the order of increasing pressure and decreasing pressure.

### Pure-component sieving coefficient measurements

The purified RNA fractions were diluted to a final volume of 10 mL and a final concentration from 15 to 18 ng/mL in tris-ethylenediaminetetraacetic acid (TE) buffer (Thermo Fisher Scientific). The ultrafiltration system was cleaned by flowing 100 mL of DEPC-treated water through the membrane. The cleaning process was stopped, and the remaining water was poured from the cell. The RNA solution was loaded into the cell immediately to prevent the membrane from drying. After stirring for 5 min, an initial sample of 100 µL was taken from the cell. The cell was closed and pressurized to the setpoint pressure. The stirring speed was 200 rpm. The system stabilized for a few minutes before the permeate line was opened. Approximately 0.5 mL of permeate, equivalent to the system hold-up volume, flowed through the permeate line before sample collection.

The sieving coefficients of each purified RNA conformer, calculated as the quotient of permeate concentration and feed concentration, were determined at different permeate fluxes. The feed concentration represented the average concentration before and after collecting the permeate sample. Concentrations were measured using Quant-it RiboGreen RNA Quantitation kits (Thermo Fisher Scientific). This highly sensitive fluorescence-based assay ranges from 1 ng/mL to 1000 ng/mL RNA. The dye was diluted 200:1 in TE buffer. One hundred microliters of sample were mixed with 100 µL of diluted dye in a well of a 96-well plate, along with calibration samples prepared by diluting known concentration solutions provided by the manufacturer. The plate was incubated in the dark for 5 min and inserted into a Synergy H1 microplate reader (Agilent, Santa Clara, CA, USA). The samples were excited at 485 nm, and the fluorescence emission intensity was measured at 530 nm. The experiment was carried out with the lights off to prevent dye photobleaching.

### Binary mixture batch diafiltration

We used the batch UF-diafiltration process in **Figure 1** to evaluate purifying circRNA from binary mixtures of the purified conformers.

**Figure 1.**
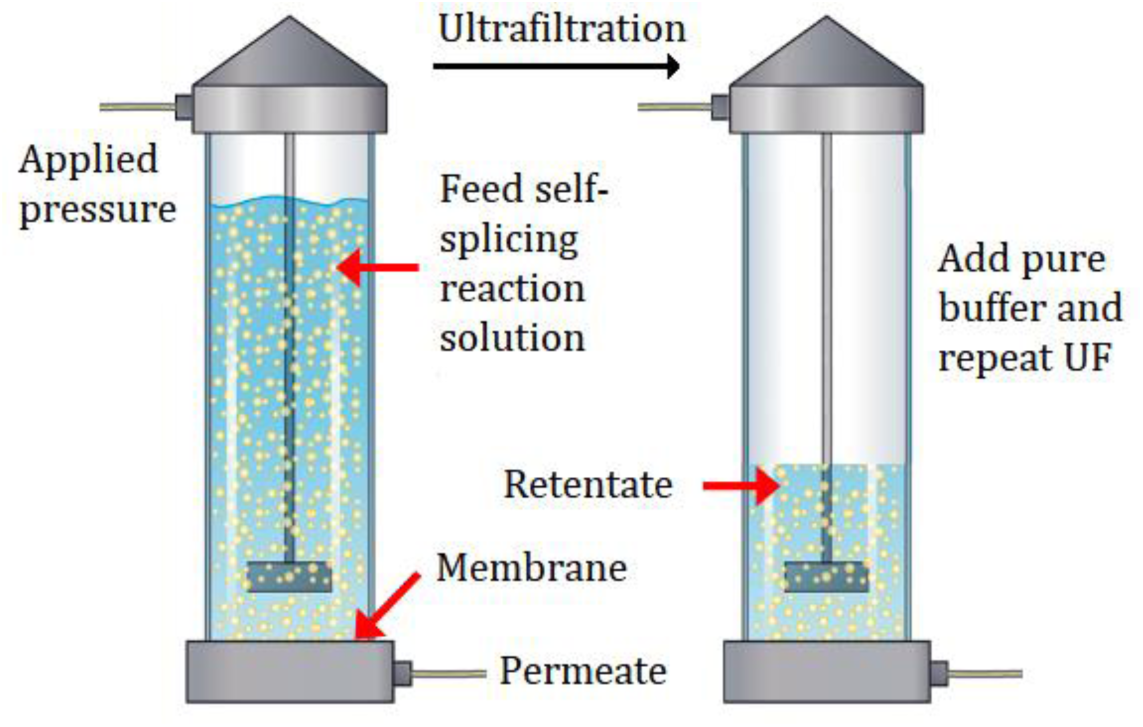
Batch direct-flow UF-diafiltration setup and process.

The first experiment used a feed solution comprising 75% circRNA and 25% preRNA. The experiment was performed three times. The percentage of circRNA in the feed varied from 74% to 77%. The second experiment used a 50:50 mixture of circRNA and nicked RNA. In each run, the initial total RNA concentration was 9.5 ng/mL, and the initial volume was 36 mL. The solution was loaded into the stirred direct-flow UF cell, pressure was applied, and half of the solution was removed as permeate, corresponding to a volume reduction factor (VRF) of 2. Pure TE buffer was added to increase the solution volume within the UF cell to its original feed volume. This cycle was performed until four diavolumes (total volume of diafiltration buffer or filtrate divided by retentate volume) were processed. Retentate samples were collected to determine circRNA purity (**Equation 1**) and yield (**Equation 2**) as a function of diavolumes.

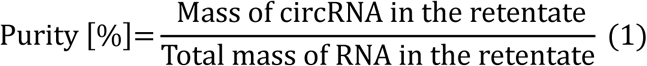

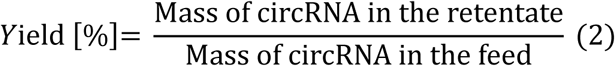

Total RNA concentration was measured using Quant-it RiboGreen RNA Quantification kits. Gel electrophoresis was used to separate the RNA conformers. The percentage of circRNA was determined by densitometric analysis using a Bio-Rad ChemiDoc XRS+ gel imaging system controlled by Image Lab 6.1 software. When the circRNA purity was higher than 90%, the linear (nicked and precursor) RNA concentrations were too low to detect by densitometric analysis. Therefore, RNase R degradation was used to calculate purity. Three samples were taken. The first was used to measure the total RNA concentration. The second was treated with RNase R by dosing one unit of RNase R per microgram of RNA, and then it was incubated at 37°C for 15 min. The third was a control sample; it was incubated as the RNase R sample at 37°C for 15 min, but without adding RNase R. An equivalent volume of TE buffer was added to keep the same volume in both samples. The control sample was used to ensure that the change in concentration was due to linear RNA degradation. The mass of circRNA was estimated by measuring the concentration of the RNase R treated sample. The total mass was measured using the concentration of the control sample. The purity was calculated by dividing the circRNA mass by the total concentration.

### Ternary mixture batch diafiltration

We evaluated purifying circRNA from a ternary mixture of the purified conformers comprising 44.4% circRNA, 23.8% preRNA RNA, and 31.8% nicked RNA. The batch diafiltration, sampling, and measurement protocols were identical to the binary mixture experiments. The purpose of this experiment is to emulate the characteristic compositions in IVT products samples to see if out protocol would work on these conditions.

### Ultrafiltration and SE-HPLC comparison

We compared separation performance by ultrafiltration and size exclusion high-performance liquid chromatography (SE-HPLC) using a mixture comprising 31.9% circRNA, 37.5% preRNA and 30.7% nicked RNA. The initial concentration was 50 ng/mL for ultrafiltration, and the initial volume was 50 mL. We ran batch diafiltration with a VRF of 2 until four diavolumes were processed. Gel electrophoresis and RNase R degradation were used to measure circRNA purity, and fluorimetry was used to measure total RNA concentration and calculate yield. For the SE-HPLC separation, the RNA mixture was pumped through a 4.6 × 300 mm size-exclusion column with a particle size of 5 µm and pore size of 200 Å (Sepax Technologies, Newark, Delaware, USA; part number: 215980P-4630) on an Agilent 1100 Series HPLC (Agilent, Santa Clara, CA) according to the protocol given by Wesselhoeft et al ^4,12^. Two HPLC runs were performed. For the first run, 30 µg of the IVT product solution was diluted in 50 µL of TE buffer and injected into the column at a 0.3 mL/min flow rate. For the second run, 19 µg of the IVT product solution was diluted in 50 µL and injected into the system at a 0.2 mL/min flow rate. Samples for each peak were collected manually. We used gel electrophoresis to determine the composition of the samples collected using a Bio-Rad ChemiDoc XRS+ gel imaging system controlled by Image Lab 6.1 software, as we did for binary and ternary mixtures. These data were used to estimate circRNA purity; the total RNA concentration was measured with fluorometry.

## Results

### circRNA synthesis and gel separation

We synthesized circular RNA using the plasmid and methods developed by Chen et al.[2, 13, 15] Gradient PCR was performed to determine that 68°C was the ideal primer annealing temperature for amplifying the NanoLuc cassette with circRNA-synIRES-R25-NanoLuc (AddGene 188116) (**Figure S2**). Purified PCR products were used as templates for RNA synthesis by IVT.

Using e-gel electrophoresis, we identified the three bands corresponding to circular, preRNA and nicked RNA (**Figure 2A**). A sample of the IVT product was treated with RNase R, which only degrades linear RNA, to confirm that we produced circRNA and to identify which e-gel band corresponded to it. Gel RNA extraction was used to obtain purified circRNA, preRNA, and nicked RNA for the ultrafiltration studies (**Figure 2B**).

**Figure 2.**
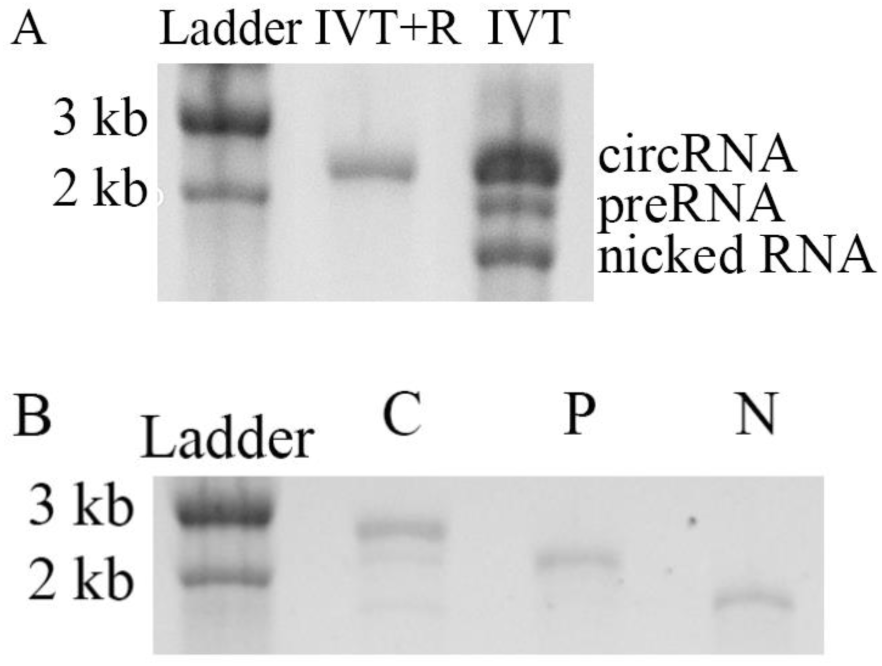
circRNA synthesis and gel purification. A) The second column represents IVT products after DNase I treatment to degrade the DNA template, the first column represents IVT products after DNase I treatment and RNase R treatment to degrade no circular RNA. B) Purified fractions of circRNA, preRNA, and nicked circular RNA were obtained using gel RNA extraction.

### RNA hydrodynamic diameter measurements

Ultrafiltration can separate nucleic acids based on size and shape[13, 15]. Before filtration, the molecule has a globular shape. As filtration begins, the converging flow field that develops near the pore entrance stretches the molecule. The molecule transitions from a globular to a fully extended shape at a high enough permeate flux. This change in shape allows the molecule to flow through the UF membrane. This point is called critical flux. An effective UF membrane allows the molecules to flow through in a fully extended shape but not in a globular shape. Therefore, to select a suitable membrane, it is essential to know the hydrodynamic diameter of the molecule in its globular shape. The nominal pore diameter of the membrane should be smaller than the hydrodynamic diameter of the circRNA.

Dynamic light scattering (DLS) is a technique that enables measurement of hydrodynamic diameter. **Table 1** shows the DLS results. The values represent the hydrodynamic diameters of the RNA in their globular state. **Table 2** reports the estimated nominal pore diameter for the membranes from Bhaumik and Santanu[16]. The RNA molecule is expected to flow through the membrane in the fully extended shape but not in the globular shape. Therefore, the membranes MWCOs selected to test had pore size smaller than all the RNA conformers dynamic diameter.

**Table 1.** DLS results for RNA conformers in TE buffer pH 7.7±0.1.

| RNA conformer | Diameter (nm) |
| --- | --- |
| circRNA | 42 |
| preRNA | 50 |
| nicked RNA | 49 |

**Table 2.** Estimated nominal membrane pore sizes from Bhaumik and Santanu[16].

| Membrane | Pore size (nm) |
| --- | --- |
| 30 kD | 11 |
| 100 kD | 12 |
| 300 kD | 20 |

### Permeability measurements

RNA samples are expensive and laborious to synthesize and purify. The smallest UF cell from the manufacturer is 50 mL. So, we designed a volume-reducing insert (**Figure S1**). We also reduced the exposed membrane area by applying an impermeable aluminum tape with holes ranging in diameter from 6 to 18 mm to decrease permeate flux. **Figure S3** shows the flow rate data collected using Biomax polyethersulfone (PES) 300 kDa membranes with different exposed areas. The permeability (slope of the lines divided by area) was the same within the measurement uncertainty (58.3 ± 0.3 L m^-2^ h^-1^ kPa^-1^) for all samples, indicating that the application of the aluminum tape ring successfully adjusted the membrane exposed area precisely. Using the smallest membrane exposed area allowed us to run the subsequent RNA ultrafiltration experiments in a time that was long enough to make accurate flow rate measurements while collecting the 100 µL sample volumes that we needed to measure concentrations.

**Figure 3** shows flux measurements using the Biomax PES 100 kDa, 50 kDa and 30 kDa membrane; the diameters were 6 mm for the 100 kDa membrane, 10 mm for the 50 kDa membrane, and 18 mm for the 30 kDa membrane. The resulting permeabilities were 2.2 µm/s/kPa for the 100 kDa membrane, 1.5 µm/s/kPa for the 50 kDa membrane, and 1.2 µm/s/kPa for the 30 kDa membrane. We demonstrated the decrease of permeability as we select lower MWCOs while operating in the same range of pressure.

**Figure 3.**
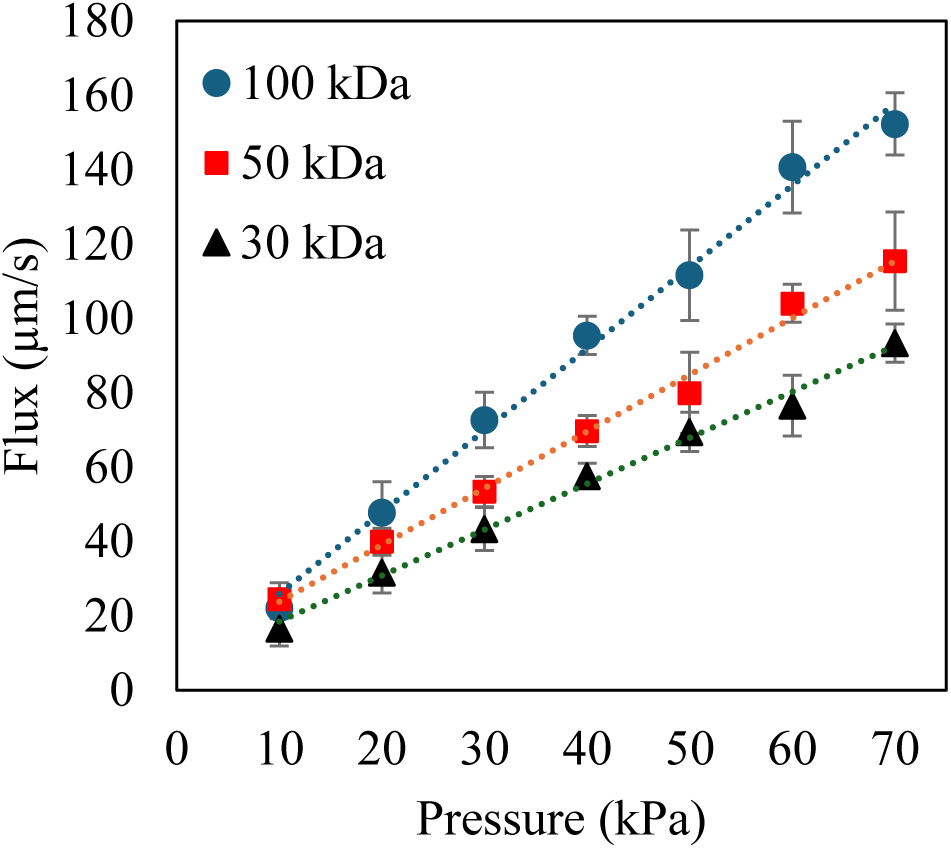
Dependence of permeability on membrane molecular weight cut off in a range from 10 to 70 kPa. As the MWCO gets higher, the permeability gets higher.

### Ultrafiltration measurements

The sieving coefficient (S) quantifies the ratio between the solute concentration in the permeate and the feed across a membrane. The solute flows freely through the membrane when S equals one. The membrane fully retains the solute when S equals zero. **Figure 4** shows the sieving coefficient measurements using purified circRNA. S (= [circRNA]_permeate_/[circRNA]_feed_) for the Biomax PES 300 kDa membrane was about 1 for all flux values, indicating that there was no retention of circRNA. Measurements with purified nicked RNA yielded the same result. The 300 kDa molecular weight cutoff (i.e., pore size) is therefore too large.

**Figure 4.**
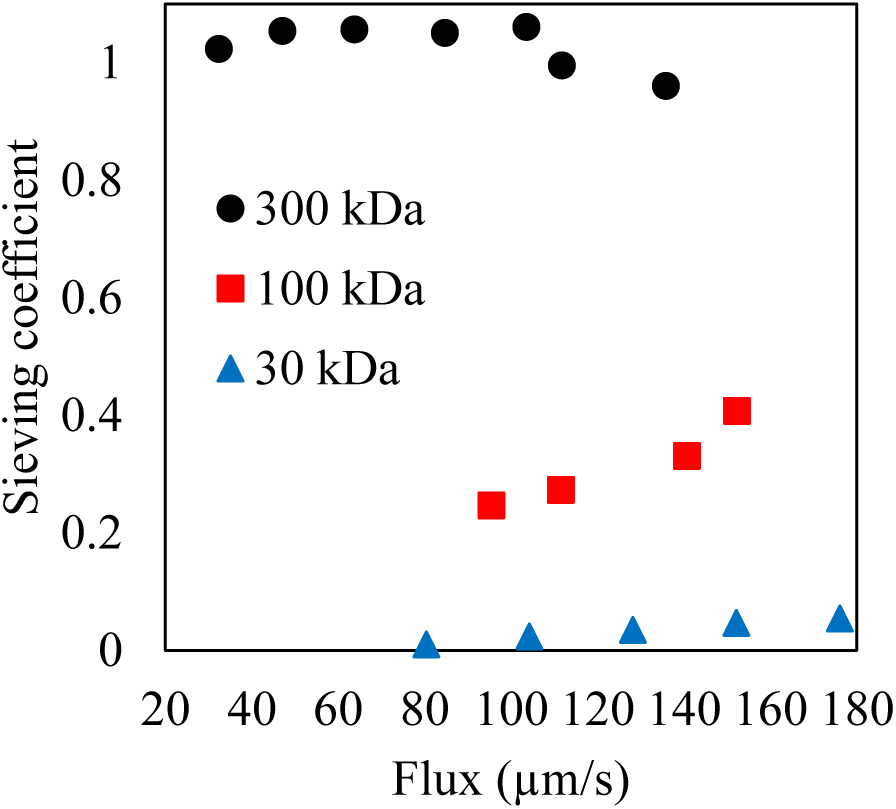
Preliminary sieving coefficients for purified circRNA using Biomax PES 300 kDa, 100 kDa and 30 kDa membranes. These data were used for membrane selection. 30 kDa membrane was selected because the permeability was close to zero.

We then measured S values for the 100 kDa membrane using purified circRNA. **Figure 4** shows the expected behavior for retention of circRNA, whereby sieving coefficients increase with increasing flux. Unfortunately, the critical flux required for transmission of circRNA, as determined by extrapolating to S = 0, gave a near zero value. Since the operating flux needs to be below the critical flux for circRNA and above that for nicked circRNA, the 100 kDa membrane also has a pore size that is too large. **Figure 4** indicates that a 30 kDa MWCO UF membrane is suitable for further testing. S for circRNA was low (below 0.06) for all flux values tested, suggesting that most circRNA was retained by the membrane at these low flux values.

The sieving coefficient results in Figure 5 indicate that a 30 kDa membrane can separate circRNA from preRNA and nicked RNA. S for circRNA was low for all flux values tested, suggesting that most circRNA was retained by the membrane. In contrast, nicked and pre-RNA sieving coefficients increased significantly for flux values higher than 200 µm/s. The data for preRNA and nicked RNA were fit to **Equation 3** with limits set between 0 and 1.

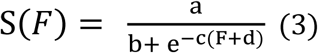

**Figure 5.**
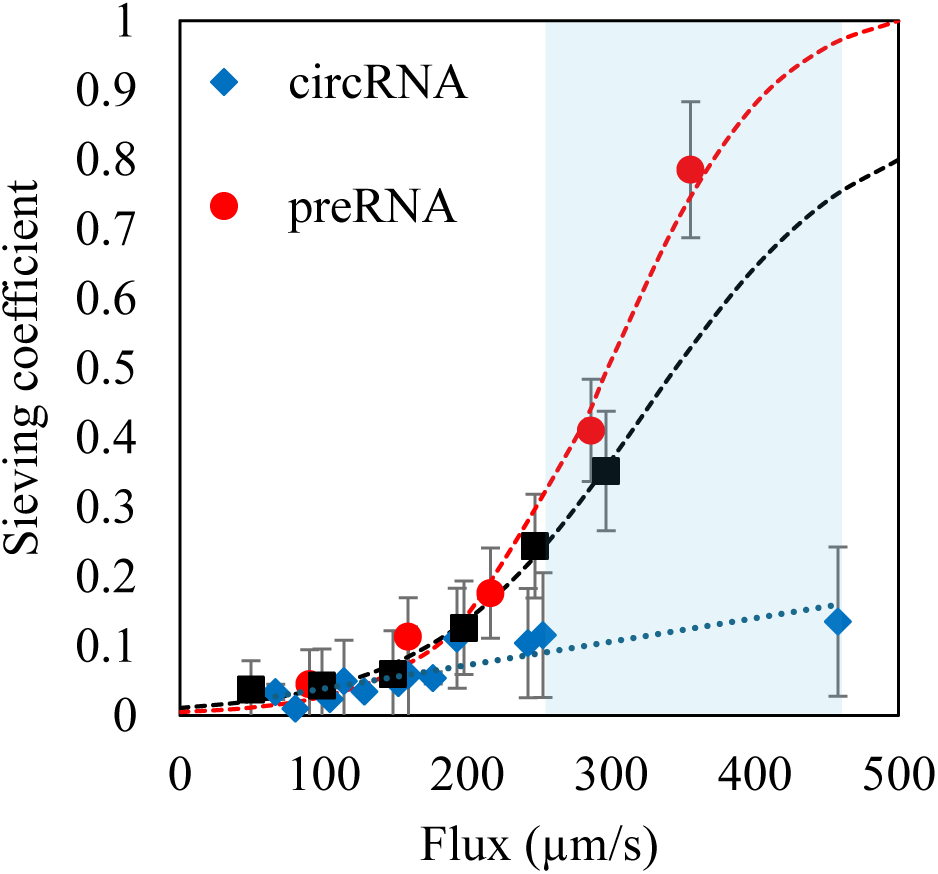
Sieving coefficient measurements for purified fractions of circRNA, nicked RNA, and preRNA using a 30 kDa Biomax membrane. The blue section represents the range of flux where we want to perform the ultrafiltration/diafiltration. In this range of conditions, we would have purity and yield above 60% for the lower flux and purity above 95% with a yield around 50% for the higher.

**Equation 3** is based on the concentration polarization model developed by Manzano and Zydney[17, 18] using a mass balance at the boundary layer.

The circRNA data were fit to an equation for a line with a slope of -1/c, which is the bulk mass transfer coefficient. The fitting parameters from Equation 3 are available in **Table 3**.

**Table 3.** Fitting parameters for purified fractions sieving coefficient model.

| preRNA data fitting |  | nicked RNA data fitting |  |
| --- | --- | --- | --- |
| a | 0.53214 | a | 0.3892 |
| b | 0.517566 | b | 0.446033 |
| c | 0.017913 | c | 0.013577 |
| d | -264 | d | -264 |

The critical flux values (S_0_) were determined from the model as the flux at which the sieving coefficient fell below 0.06. This value is equal to the uncertainty in the sieving coefficients, derived from propagating the uncertainties in the concentration measurements. The critical flux for circRNA is 160 µm/s, for preRNA is 144 µm/s, and for nicked RNA is 134 µm/s. Although the difference among critical fluxes is small, the sieving coefficient of circRNA is significantly lower than the sieving coefficients of preRNA and nicked RNA at higher flux values. The low, near-constant sieving coefficient for circRNA may be due more to the distribution of pore sizes in the membrane than flow-induced deformation, whereby a small fraction of circRNA molecules is transmitted through the larger pores in the distribution. This differs from the linear molecules, where the large increase in transmission at higher flux is likely due primarily to elongation in the converging flow field near pore entrances.

### Binary purification measurements

**Figure 6** shows diafiltration results for a mixture of circRNA and preRNA. The points represent experimental data, and the curves represent the model predictions using the sieving coefficients data measured for the purified fractions. Equations 4 and 5 were used to calculate circRNA yield (Y_circ_) and purity (P_circ_).

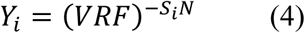

**Figure 6.**
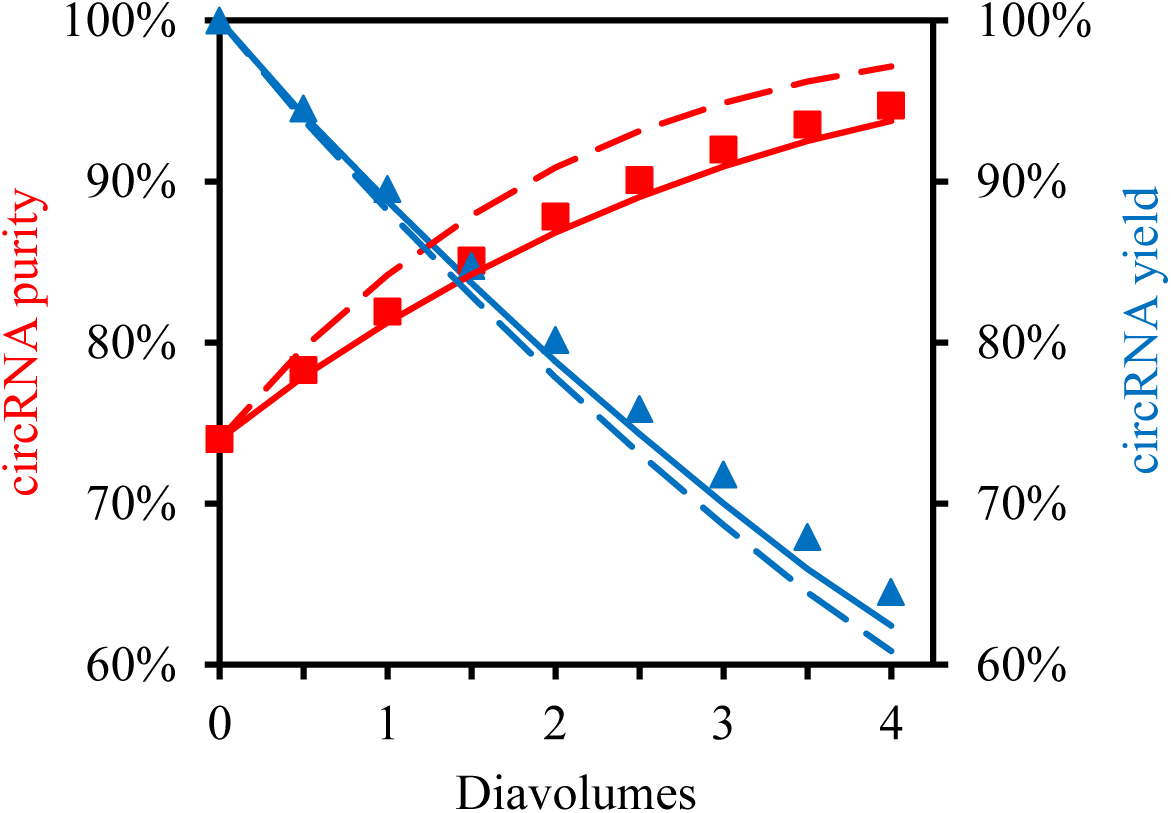
Diafiltration of a mixture of circRNA and preRNA. Experimental data shown as symbols were measured at a flux between 310 and 274 µm/s. The dashed curve represents the model predictions using a constant flux of 310 µm/s, and the continuous curve represents the model predictions using a constant flux of 274 µm/s.

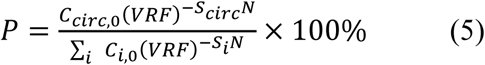

N indicates the number of cycles. C_i,0_ is the initial concentration of component i. A circRNA purity of 95% and a yield of 62% were achieved after 3.9 diavolumes. The permeate flux during the experiment decreased from 310 to 274 µm/s while keeping a constant pressure of 414 kPa. The figure shows two model curves: the dashed curve represents the model prediction using sieving coefficients at the highest flux, and the continuous curve represents the model prediction using sieving coefficients at the lowest flux. The experimental results align well with the model predictions.

**Figure 7** shows diafiltration results for a mixture of circRNA and nicked RNA. We achieved a circRNA purity of 80% with a yield of 50%. More significant deviations were observed between the model predictions and the experimental data for this mixture. The flux decreased from 355 to 272 µm/s during the experiment. Therefore, the effect of flux decline on separation selectivity was more significant. Again, the dashed curve represents the model predictions at the highest flow, and the continuous curve represents the model predictions at the lowest flux. The circRNA experimental purity was closer to the prediction at the lowest flux. However, the yield was lower than we would expect at that flux. A more significant decrease in the flux during the experiment may be related to fouling issues.

**Figure 7.**
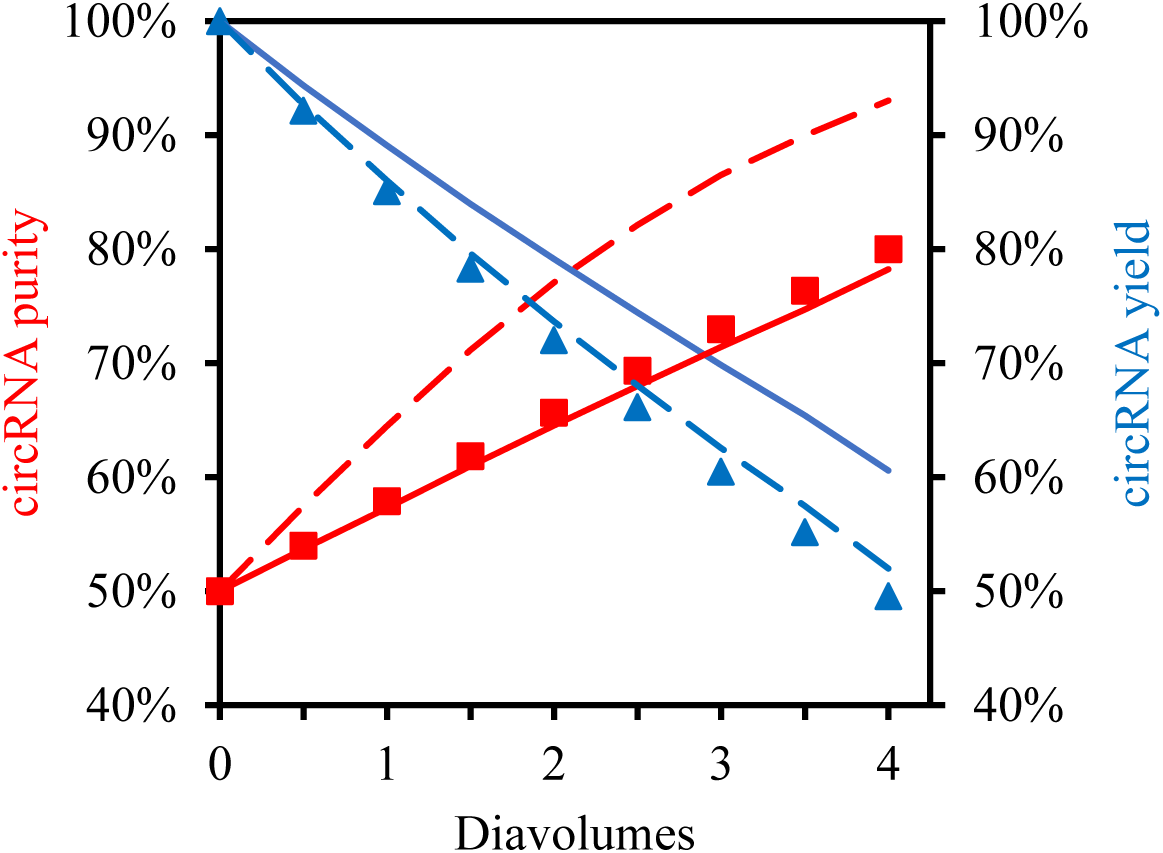
Diafiltration of a binary mixture of circRNA and nicked RNA. Experimental data were measured at a flux between 272 and 355 µm/s. The dashed curve represents the model predictions using a constant flux of 355 µm/s, and the continuous curve represents the model predictions using a constant flux of 272 µm/s.

### Ternary mixture measurements

**Figure 8** shows diafiltration results for the ternary mixture. We achieved a circRNA purity of 96% with a yield of 53% after four diavolumes. The flux dropped from 425 to 350 µm/s. As before, the dashed curve represents the model predictions at the highest flux, and the continuous curve represents predictions at the lowest flux. The experimental results align well with the model predictions.

**Figure 8.**
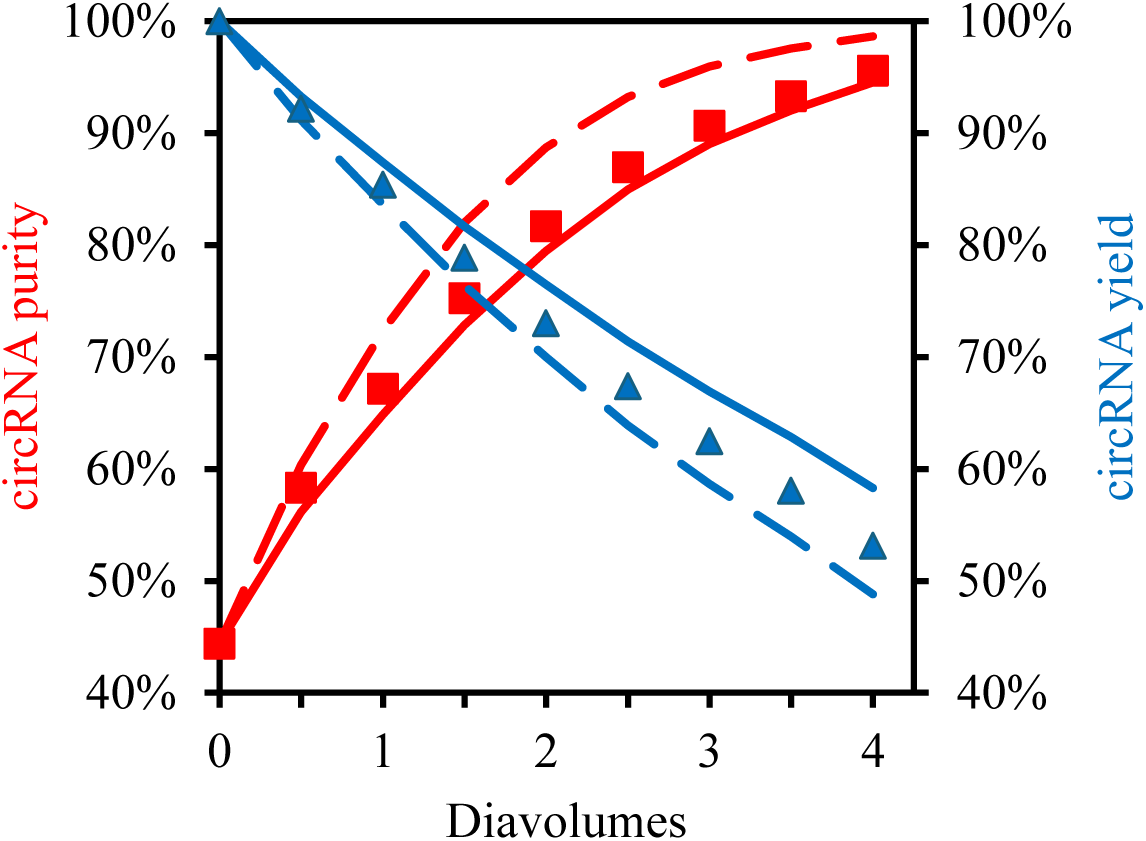
Diafiltration of a ternary mixture of circRNA (44.4%), preRNA (23.8%), and nicked RNA (31.8%) at a flux above the critical flux of preRNA and nicked RNA. The dashed curve represents the model predictions using a constant flux of 425 µm/s, and the continuous curve represents the model predictions using a constant flux of 350 µm/s.

### IVT products ultrafiltration and SE-HPLC comparison

**Figure 9** shows diafiltration results for the IVT solution. We achieved a circRNA purity of 81% with a yield of 65% after four diavolumes. The flux dropped from 334 to 258 µm/s. As before, the dashed curves represent the model predictions at the highest flux, and the continuous curves represent predictions at the lowest flux. The experimental results align well with the model predictions. The purity achieved was not as high as that obtained for ternary mixtures. This may be related to impurities such as introns, and residual nucleotides and DNA template.

**Figure 9.**
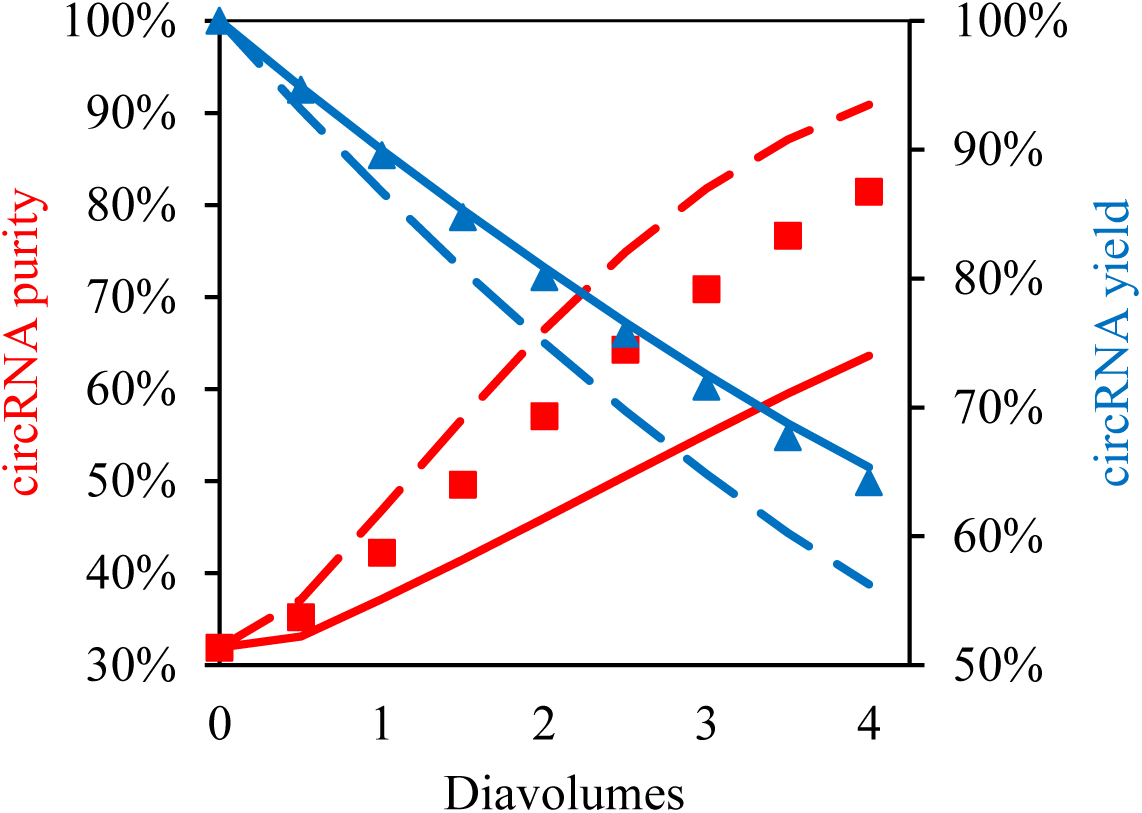
Diafiltration of a diluted IVT solution containing circRNA (31.9%), preRNA (37.5 %), and nicked RNA (30.7%) at a flux above the critical flux of preRNA and nicked RNA yielded a purity of 81% with a yield of 61%.

**Figure 10** shows the SE-HPLC chromatogram for the same IVT products used for the ultrafiltration experiment. The red area indicates the sample collected. The purity of the sample collected was measured using a densitometric analysis, and the total RNA concentration was measured using fluorometry. The purity was 41%, and the yield was 45%. It is possible to obtain higher purity by collecting a narrower peak. As you can see in **Table 4**, when collecting a narrower sample, the yield drastically decrease. Therefore, we conclude that at this conditions it is not possible to improve purity without sacrificing yield.

**Figure 10.**
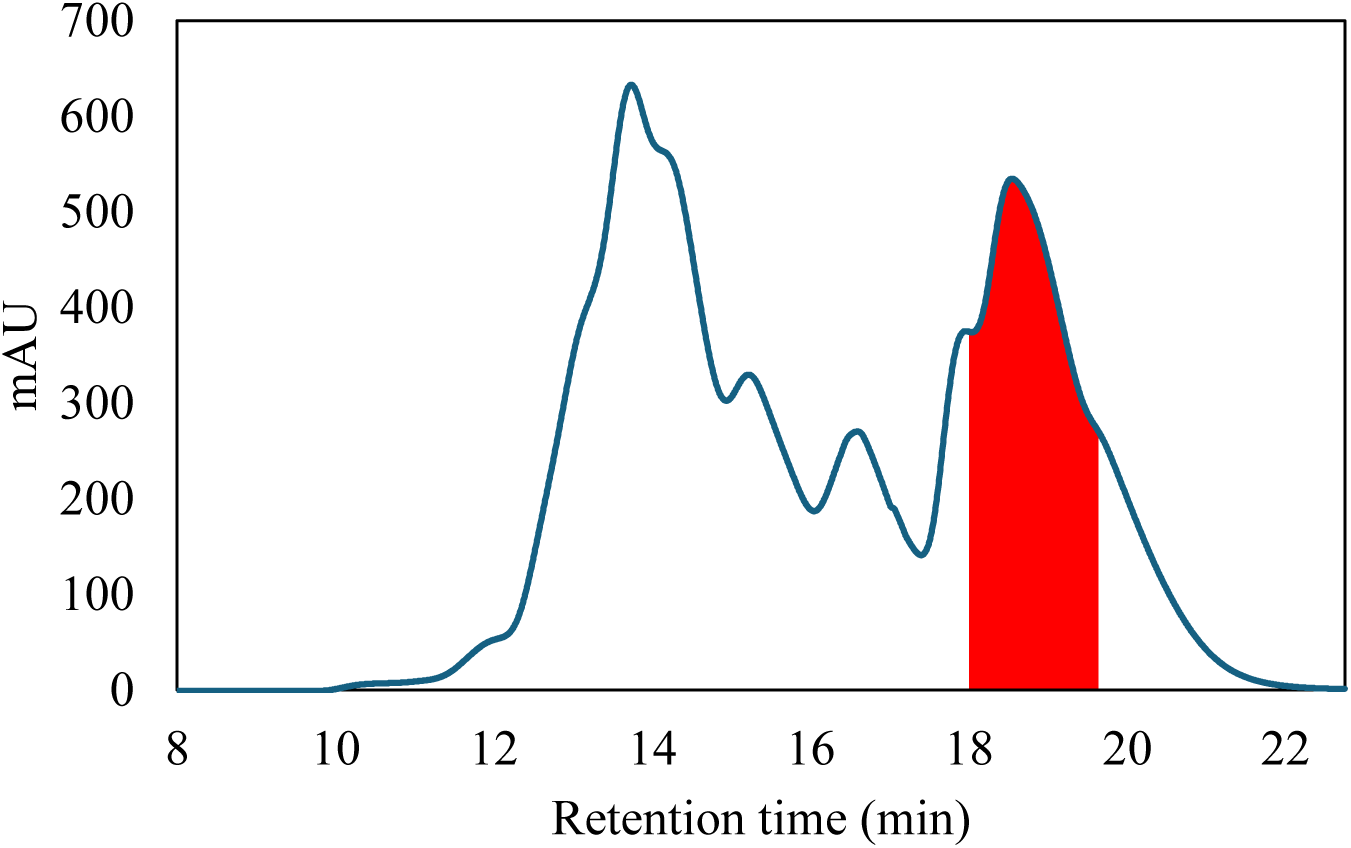
SE-HPLC purification of a diluted IVT solution containing circRNA (31.9%), preRNA (37.5 %), and nicked RNA (30.7%).

**Table 4.**
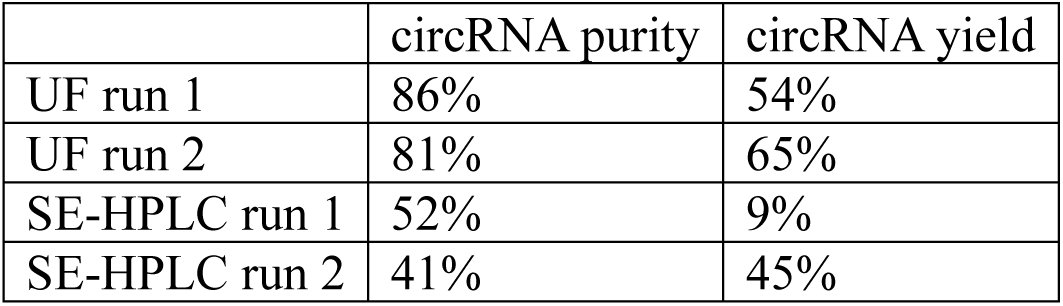
Summary of circRNA purity and yield for UF and SE-HPLC runs using IVT solutions.

|  | circRNA purity | circRNA yield |
| --- | --- | --- |
| UF run 1 | 86% | 54% |
| UF run 2 | 81% | 65% |
| SE-HPLC run 1 | 52% | 9% |
| SE-HPLC run 2 | 41% | 45% |

**Table 4** summarizes the results we obtained for the purification of circRNA from IVT solutions using UF and SE-HPLC. UF experiments were carried out at the same pressure of 414 kPa. However, the fluxes were different due to slight differences in membrane permeability. Nevertheless, we obtained high circRNA purity and yield for both UF diafiltration experiments. Obtaining high-purity circRNA from SE-HPLC was challenging due to overlapping peaks. Both purification methods observed a tradeoff between circRNA purity and yield. However, purity and yield were dramatically lower for SE-HPLC than UF.

### Conclusions

RNA therapies have the potential to revolutionize medicine, but challenges in ensuring RNA stability and purity remain. Circularizing RNA increases its stability, though current methods for purifying circRNA, like SE-HPLC combined with RNase R digestion, have limitations in purity and yield. Our attempts to replicate these methods achieved only 41% purity and 45% yield. An attempt to increase purity by selecting a narrow peak led to a drastically lower yield. This highlights the need for new, scalable purification techniques to achieve high purity and yield.

We explored ultrafiltration as an alternative for purifying circRNA and achieved higher purity and yield than SE-HPLC. The purity is around double the SE-HPLC achieved, and the yield is above 50%. Additionally, the relative ease of scaling the ultrafiltration process increases its potential for purifying commercial circRNA therapeutics. A challenge to overcome is the variation of sieving coefficients during operation due to flux decline associated with membrane fouling and concentration polarization. The next step is to demonstrate the ability of circRNA purified by ultrafiltration to express a gene in mammalian cells and *in vivo*. The results of this study may improve the affordability and accessibility of RNA-based medical treatments and vaccines.

## Author Contributions

Conceptualization, K.G., M.R.B and S.M.H.; methodology, K.G., X.L., M.R.B. and S.M.H.; validation, K.G., X.L., M.R.B. and S.M.H.; formal analysis, K.G., X.L., M.R.B. and S.M.H.; investigation, K.G. and X.L.; resources, M.R.B. and S.M.H.; data curation, K.G. and X.L.; writing—original draft preparation, K.G. and X.L.; writing—review and editing, M.R.B. and S.M.H.; visualization, K.G.; supervision, M.R.B. and S.M.H.; project administration, S.M.H.; funding acquisition, M.R.B. and S.M.H. All authors have read and agreed to the published version of the manuscript.

## Data Availability Statement

The data required to generate the figures can be obtained from https://github.com/kguille19/Purifying-circular-RNA-from-in-vitro-transcription-reactions-by-ultrafiltration. Additional data may be provided on request.

## Acknowledgments

S.M.H. acknowledges support from the William B. “Bill” Sturgis, ‘57 and Martha Elizabeth “Martha Beth” Blackmon Sturgis Distinguished Professorship in Chemical and Biomolecular Engineering.

## References

1. Li, H., et al., Circular RNA cancer vaccines drive immunity in hard-to-treat malignancies. 2022. p. 6422–6436.

2. Chen, R., et al., Engineering circular RNA for enhanced protein production. 2022, Springer US.

3. Wesselhoeft, R.A., P.S. Kowalski, and D.G. Anderson, Engineering circular RNA for potent and stable translation in eukaryotic cells. 2018. p. 2629.

4. Cheng, J., et al., Circular RNAs with protein-coding ability in oncogenesis. 2023, Elsevier B.V. p. 188909.

5. Yuan, W., X. Zhang, and H. Cong, Advances in the protein-encoding functions of circular RNAs associated with cancer (Review). 2023. p. 1–14.

6. Kameda, S., H. Ohno, and H. Saito, Synthetic circular RNA switches and circuits that control protein expression in mammalian cells. 2023, Oxford University Press. p. E24.

7. Tai, J. and Y.G. Chen, Differences in the immunogenicity of engineered circular RNAs. 2023.

8. Niu, D., Circular RNA vaccine in disease prevention and treatment. 2023, Springer US.

9. Kristensen, L.S., et al., The biogenesis, biology and characterization of circular RNAs. 2019, Springer US. p. 675–691.

10. Feng, X., et al., Messenger RNA chromatographic purification: advances and challenges. Journal of Chromatography A, 2023. 1707: p. 464321.

11. Abe, B.T., et al., Circular RNA migration in agarose gel electrophoresis. 2022, The Authors. p. 1768–1777.e3.

12. Wesselhoeft, R.A., et al., RNA Circularization Diminishes Immunogenicity and Can Extend Translation Duration In Vivo. 2019. p. 508–520.e4.

13. Latulippe, D.R. and A.L. Zydney, Separation of plasmid DNA isoforms by highly converging flow through small membrane pores. 2011. p. 548–553.

14. Chen, R., et al., Engineering circular RNA for enhanced protein production. 2022, Springer US.

15. Latulippe, D.R. and A.L. Zydney, Elongational flow model for transmission of supercoiled plasmid DNA during membrane ultrafiltration. 2009. p. 201–208.

16. Sutariya, B. and S. Karan, A realistic approach for determining the pore size distribution of nanofiltration membrane*s*. 2022, Elsevier B.V. p. 121096.

17. Latulippe, D.R., K. Ager, and A.L. Zydney, Flux-dependent transmission of supercoiled plasmid DNA through ultrafiltration membranes. 2007. p. 169–177.

18. Manzano, I. and A.L. Zydney, Quantitative study of RNA transmission through ultrafiltration membranes. 2017, Elsevier B.V. p. 272–277.

