## Supplementary material for "Purifying circular RNA by ultrafiltration"

Supplementayr information

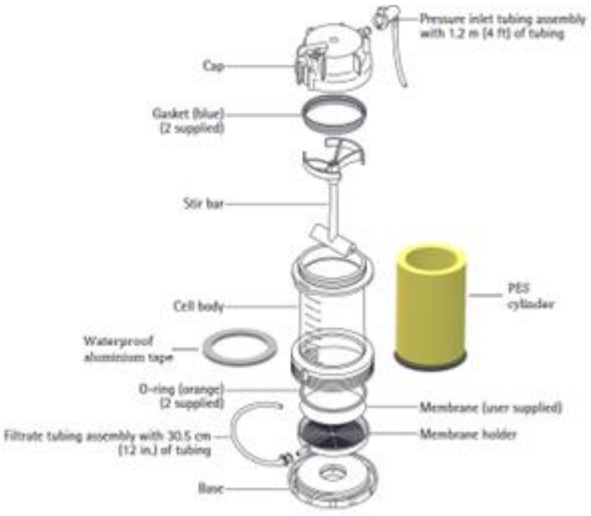


**Figure S1.** UF cell modification to decrease volume and flow. Ultrafiltration cell diagram including design modifications for this project.

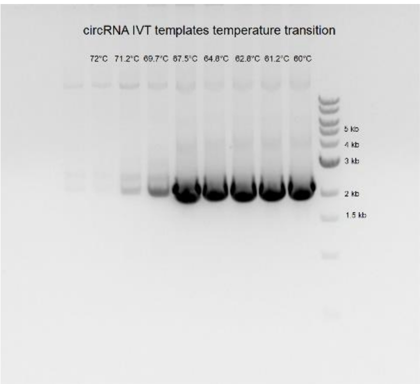


**Figure S2.** Gradient primer annealing temperature for PCR.

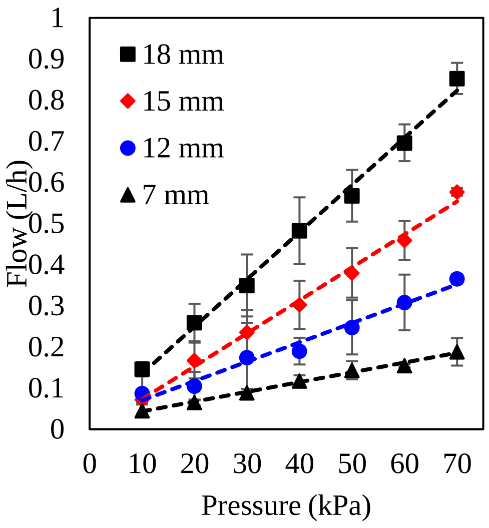


**Figure S3**. Dependences of pressure and exposed membrane diameter on flow rate.

**Table S3.** PCR protocol provided by New England Biolabs**.** The template was amplified using the primers provided by Chen et al. A) Temperature gradient. B) Amplification.

A

| | | |
|---|---|---|
| Initial Denaturation | 98°C | 30 s |
| 30 Cycles | 98°C<br>60-72°C<br>72°C | 10 s<br>30 s<br>1 min |
| Final Extension | 72°C | 2 min |
| Hold | 4–10°C | ∞ |

B

| | | |
|---|---|---|
| Initial Denaturation | 98°C | 30 s |
| 30 Cycles | 98°C<br>68°C<br>72°C | 10 s<br>30 s<br>1 min |
| Final Extension | 72°C | 2 min |
| Hold | 4–10°C | ∞ |
